# Effects of a 33-ion sequential beam galactic cosmic ray analog on male mouse behavior and evaluation of CDDO-EA as a radiation countermeasure

**DOI:** 10.1101/2021.07.15.451917

**Authors:** Frederico C Kiffer, Krishna Luitel, Fionya H Tran, Riya A Patel, Catalina S Guzman, Ivan Soler, Rui Xiao, Jerry W Shay, Sanghee Yun, Amelia J Eisch

## Abstract

In long-term spaceflight, astronauts will face unique cognitive loads and social challenges which will be complicated by communication delays with Earth. It is important to understand the central nervous system (CNS) effects of deep spaceflight and the associated unavoidable exposure to galactic cosmic radiation (GCR). Rodent studies show single- or simple-particle combination exposure alters CNS endpoints, including hippocampal-dependent behavior. An even better Earth-based simulation of GCR is now available, consisting of a 33-beam (33-GCR) exposure. However, the effect of whole-body 33-GCR exposure on rodent behavior is unknown, and no 33-GCR CNS countermeasures have been tested. Here astronaut-age-equivalent (6mo-old) C57BL/6J male mice were exposed to 33-GCR (75cGy, a Mars mission dose). Pre-/during/post-Sham or 33-GCR exposure, mice received a diet containing a ‘vehicle’ formulation alone or with the antioxidant/anti-inflammatory compound CDDO‐EA as a potential countermeasure. Behavioral testing beginning 4mo post-irradiation suggested radiation and diet did not affect measures of exploration/anxiety-like behaviors (open field, elevated plus maze) or recognition of a novel object. However, in 3-Chamber Social Interaction (3-CSI), CDDO-EA/33-GCR mice failed to spend more time exploring a holder containing a novel mouse vs. a novel object (empty holder), suggesting sociability deficits. Also, Vehicle/33-GCR and CDDO-EA/Sham mice failed to discriminate between a novel stranger vs. familiarized stranger mouse, suggesting blunted preference for social novelty. CDDO-EA given pre-/during/post-irradiation did not attenuate the 33-GCR-induced blunting of preference for social novelty. Future elucidation of the mechanisms underlying 33-GCR-induced blunting of preference for social novelty will improve risk analysis for astronauts which may in-turn improve countermeasures.

## 1. INTRODUCTION

Understanding the biological effects of spaceflight beyond low-Earth orbit remains a pressing priority in order to achieve a successful crewed mission to Mars. A major obstacle to deep spaceflight is the space radiation environment. The interplanetary radiation field consists of energetic charged-particles capable of breaching conventional spacecraft shielding. Charged-particle radiation includes low energy but high abundance particles such as protons to highly-charged, heavier but lower abundance particles such as ^16^O, ^28^Si, and ^56^Fe. These particles originate from periodic but largely unpredictable solar ejecta and from galactic cosmic radiation (GCR), the remnants of supernovae that circulate through the galaxy at a constant fluence. GCR contributes to the vast majority of the total predictable Mars mission dosage [1]. Most space missions to date have occurred within Earth’s magnetosphere, and thus astronauts have largely been shielded from these potentially damaging high-energy, charged particles. As such, there are enormous gaps in knowledge about the human consequences of GCR exposure [2]. Such knowledge is essential to prepare NASA for successful deep spaceflight missions, but will also help advance science and medicine in other fields that use charged particles, such as in particle radiation used for cancer therapy [3].

Of NASA’s many risk-based areas, the central nervous system (CNS) is of great concern due to the cognitive demands on astronauts and the social challenges associated with long-term spaceflight. These challenges are exacerbated by the lag in communication associated with increasing distances from Earth leading to lack of immediate ground-based mission support, long-term confinement, and the added psychological stress of no-emergency-return contingencies on a mission to Mars. Thus, learning, memory, and social cognition are among the most important neurocognitive domains to evaluate for their sensitivity to GCR. As modeling GCR on Earth has previously been challenging, our understanding of how CNS function is influenced by space radiation comes primarily from studies where rodents are exposed to single charged particles in NASA’s Space Radiation Laboratory (NSRL) and similar particle accelerator facilities. These single-particle studies show space radiation often [but not always; e.g. 4,5–7] diminishes aspects of rodent cognitive behavior, including memory for objects or fear, working memory, spatial navigation, attention, anxiety-like behavior, sociability, and preference for social novelty [8]. A major limitation to these studies is the use of monoenergetic, single-ion beams which are straightforward for accelerators to produce on Earth but fail to mimic the complexity of particles experienced in space. An important advancement in the recapitulation of the radiation field encountered in space is illustrated by recent work examining the impact of a simple mix of varying (2-6) charged particles on rodent behavior [8–11]. When used at Mars-relevant doses [<1.5 Gy, 1], whole-body exposure of male rodents to simple mixed beams also decreases aspects of rodent cognitive behavior. Mixed beams impair male rodent sociability, preference for social novelty, hippocampal-dependent behavior (novel object recognition [NOR], water maze) and increase measures relevant to anxiety (elevated plus maze [EPM]), with normal presentation of other behaviors (exploratory locomotion, passive avoidance, forced swim test). While studies using the simple mix of particles are seminal, this exposure paradigm still falls far short of mimicking the GCR that will be experienced during deep space travel [12]. In a major step forward for the field, recent NSRL upgrades provide an even more Mars-relevant exposure with a standardized diversity of particles and energy spectra, termed a 33-ion simulated Galactic Cosmic Ray radiation field (33-GCR)[12]. Although 33-GCR is now available, it is currently unknown how exposure to such a complex, mixed radiation field influences the CNS specifically in the context of rodent behavior.

Given that single- and simple mixed-particle studies of space radiation suggest it damages the rodent CNS [c.f. 4,5–7], it is reasonable to consider countermeasures to protect the CNS from the charged particle environment of space. Physical countermeasures, such as shielding the spacecraft or spacesuit, currently are cost prohibitive and accompanied by complications [13]. Alternative radiation mitigation strategies, such as pharmacological countermeasures or interventions, are an area of active research [13]. Since radiation exposure in space is constant, the best pharmaceutical countermeasures must also be compatible with the quality of life of a space crew. To this end, topical or oral administration of pharmacological countermeasures with long biological half-lives may be preferred. One such compound with a long biological half-life is the triterpenoid 2-cyano-3, 12-dioxooleana-1, 9-dien-28-oic acid (CDDO)-ethylamide (CDDO-EA), which targets the Nrf-2/ARE pathway, upregulates endogenous oxidative stress response elements, and potentially quenches the dense oxidative stress produced by particle radiation [14]. CDDO-EA is an effective countermeasure against ^1^H+^4^He+^28^Si-induced blood plasma lipid peroxidation and lung cancer initiation after an acute, 30cGy, whole-body dose [15]. In the CNS, CDDO-EA and structurally similar variants (such as CDDO-Imidazole or CDDO-Methyl) improve pathological symptoms in mouse models of Huntington’s disease, ischemic injury, cerebral malaria, and amyotrophic lateral sclerosis via their anti-inflammatory/antioxidant actions [14,16–19]. CDDO-Methyl is currently in Phase 3 of antioxidant-based clinical trials (NCT03749447). Although space radiation has long been known to increase CNS indices of inflammation and reactive oxygen species [20–23], no work has yet examined the ability of any CDDO variant to act as an effective CNS radiation countermeasure in rodent models of space radiation exposure.

To understand the effects of GCR on the CNS and evaluate the potential behavioral neuroprotective properties of CDDO-EA, we exposed 6-month(mo)-old male mice to sham radiation or an acute whole-body 75cGy dose of 33-GCR with or without transient co-administration of CDDO-EA. Four months later, we began behaviorally testing the mice in mission-relevant tests spanning cognitive and social domains. We hypothesized that whole-body 33-GCR exposure would decrease sociability, preference for social novelty, and hippocampal-dependent memory (such as NOR), and increase measures relevant to anxiety (such as EPM) as has been shown after exposure to simple and mixed particles. The data show that mature male mice exposed to 33-GCR normal levels of locomotion, sociability, and even – in contrast to our hypothesis – normal levels of anxiety-like behavior and hippocampal-dependent memory. While sociability was normal in most groups, mice given the combination of CDDO-EA and 33-GCR had sociability deficits. Also, mature male mice exposed to either 33-GCR or CDDO-EA had blunted preference for social novelty. Finally, CDDO-EA (given at the time of irradiation) did not block the 33-GCR-induced blunting of preference for social novelty. While our CDDO-EA countermeasure results merit additional study with different administration and radiation parameters, our finding that 33-GCR blunts preference for social novelty underscores why the CNS should remain a critical area of concern in regard to the risks of deep space flight and extraterrestrial environments.

## 2. METHODS

### 2.1 Animals

6mo-old male C57BL/6J mice (Jackson Laboratory stock #000664, weanling mates, Bar Harbor, Maine) were shipped directly to Brookhaven National Laboratories (BNL). Mice were housed 4 per cage under a regular 12:12 hour (h) light cycle at 22°C, 30-70% humidity and given standard rodent chow (LabDiet 5015 #0001328) and water *ad libitum*. After 3 days of acclimation, mice were split into 2 diet groups to receive either a vehicle (Veh: Purina Rodent Diet 5002, 12.5g EtOH, 37.5g Neobee Oil) or a CDDO-EA (Veh + 400mg/kg RTA 405; Reata Pharmaceuticals, Irvine, TX) chow *ad libitum* for 5 days. Both the Veh and CDDO-EA formulations were prepared by Purina Mills, LLC. On day 4 of Veh or CDDO-EA diet, mice in both groups were further subdivided into Sham or 33-GCR groups (*n* = 22-24 per Diet/Radiation group; **Supplemental [Supp.] Fig. 1**), as detailed in Sections 2.2 and 2.3. The day after irradiation, mice were shipped to the Children’s Hospital of Philadelphia (CHOP) via ground transport. As part of standard CHOP quarantine procedure, mice were fed 4% fenbendazole chow for 1.5 mo and then returned to standard rodent chow (LabDiet 5015 #0001328). For the remainder of the experiment, mice were housed in HEPA-filtered, closed airflow vivarium racks (Lab Products Inc., Enviro-Gard™ III, Seaford, DE) under a 12:12h light/dark cycle at 20-23°C and 30-40% humidity. Mice were weighed periodically, with one weigh bucket per cage to prevent exposure to odors from other cages. After delivery to CHOP, mice that necessitated single housing due to aggression were excluded from behavioral experiments (**Supp. Fig. 1**). All care and procedures were approved by the Institutional Animal Care and Use Committees (IACUC) at BNL and CHOP and were in accordance with the AAALAC and National Institute of Health (NIH) guidelines for the care and use of laboratory animals. Our scientific reporting adheres to the ARRIVE 2.0 guidelines [24].

### 2.2 Radiation

On the day of irradiation at the appointed time, Sham and GCR mice were placed in small well-ventilated holders (10 × 10 × 4.5 cm) paired with cagemates. GCR mice were then given an acute 75cGy whole-body exposure of the NSRL 33-beam GCR simulation over the duration of 1.5h. Radiation beam parameters and delivery order can be found in **Supp. Table 2**. Sham-irradiated mice did not receive charged-particle radiation but remained in holders for an equivalent period of time. Radiation was delivered in an even 60 x 60cm beam distribution in the Spring 2019 (19A) campaign. Dosimetry and beam calibration were provided by NSRL staff.

### 2.3 Overview of Behavioral Testing

Behavioral testing began 4 mo after irradiation and continued for ~6 mo to assess mid- to late effects (**Fig. 1A**). Testing was conducted during the light cycle under dim red light conditions (~30-50 lux) at 72°F and 35-50% humidity. Mice were acclimated to behavior rooms for 1h prior to testing. Recording for most tests was acquired by a ceiling-mounted camera (Ace acA640-90gc, Basler) and tracking was extrapolated with Ethovision XT (Noldus Information Technology). Nose points were used for exploratory measures, and center points for gross locomotor measures. Behavioral tracking for activity chambers was acquired by infrared beam sensors and processed by Activity Monitoring 5 (Med Associates Inc., #SOF-811). Once mice were placed in testing arenas, the handler left the noise-isolated behavior room and monitored mouse activity from a computer in an adjacent room. All behavioral equipment was disinfected and deodorized with 10% TB-10 (Birex) in between testing trials and allowed time to dry. Mouse handlers were blinded to experimental conditions. Sample sizes for behavioral tests are *n* = 11-15 per group. A sole Veh/33-GCR mouse was lost ~6 mo prior to the NOR test. This mouse’s behavioral data collected near the time of death was retroactively removed from two trials of the 3-Chamber Social Interaction (3-CSI; Sociability and Preference for Social Novelty trials, detailed in Section 2.7). Five additional mice (CDDO-EA/Sham n=1, CDDO-EA/33-GCR n=4) had their data removed from the 3-CSI Preference for Social Novelty trial due to equipment issues and outlier analysis (**Supp. Table 1**).

**Figure 1.**
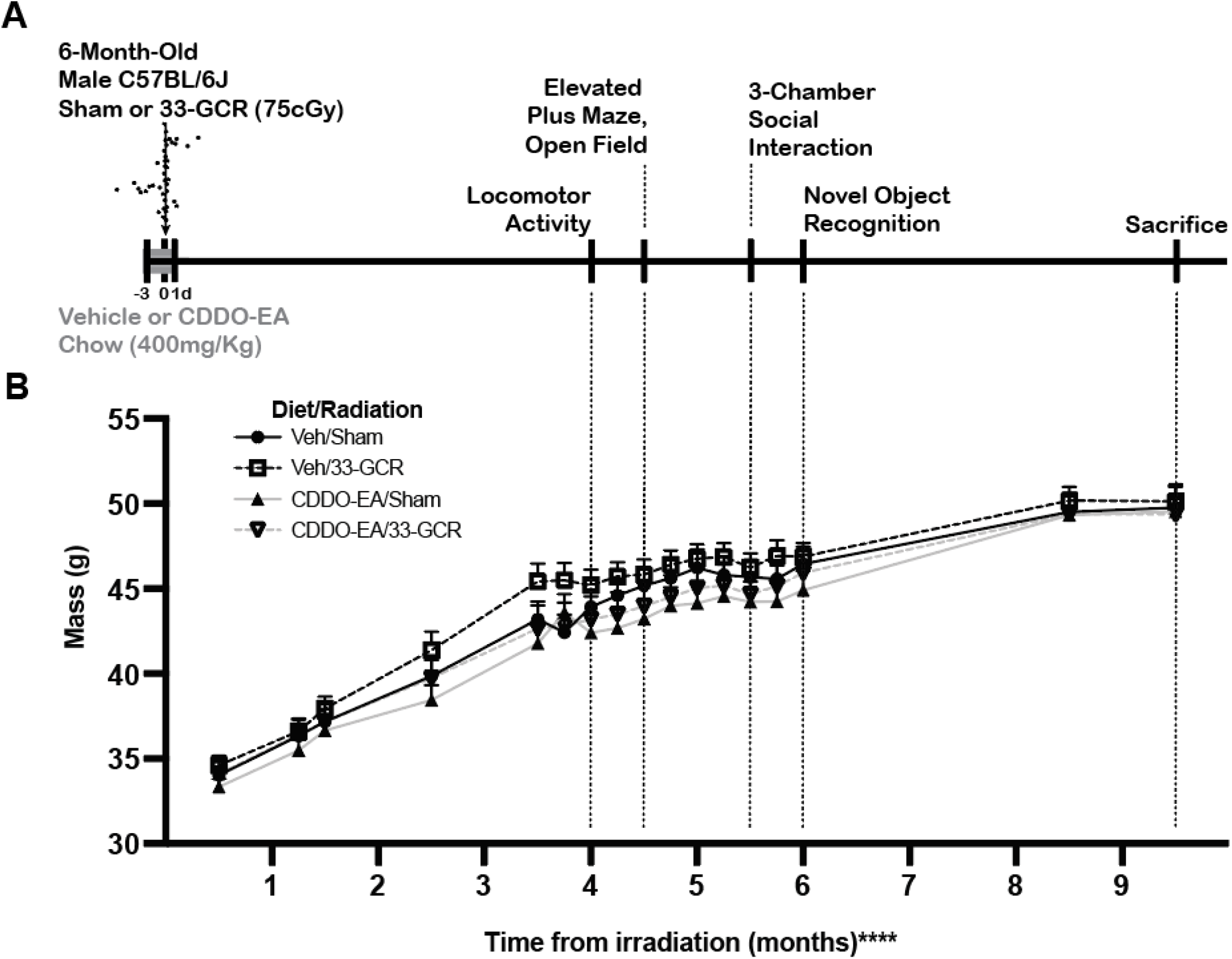
Experimental timeline and weights of behaviorally-tested mice. **A**) **Timeline**. Six-month-old male C57BL/6J mice received either a Mars-mission-relevant 75cGy whole-body exposure to ground-based galactic cosmic radiation consisting of 33 unique particle types of varying “Z” (atomic weight) and energies (33-GCR, represented by the arrow with a particle track) or sham irradiation (Sham). A subset of each group was given the candidate dietary countermeasure 2-cyano-3, 12-dioxooleana-1, 9-dien-28-oic acid-ethylamide (CDDO-EA) or a vehicle (Veh) diet for 5 consecutive days: before, during, and immediately after irradiation (indicated by gray portion on timeline). Thus, Day 4 of CDDO-EA or Veh coincided with exposure to 33-GCR or Sham. Beginning 4 months post-irradiation, mice were tested for behaviors in the order as shown. These tests were used to assess gross locomotor and exploratory activity (Locomotor Activity, Open Field, 3-Chamber Social Interaction [3-CSI] Habituation trial), anxiety-like behavior (Elevated Plus Maze, Open Field), sociability (3-CSI Sociability trial), preference for social novelty and social memory (3-CSI Preference for Social Novelty trial), and exploratory cognitive behavior (NOR). Sacrifice occurred 39 weeks post-irradiation. **B**) Body mass in all groups increased over time with no difference among groups. Dotted lines extend from timeline (**A**) to body mass data points (**B**) to facilitate understanding what was happening to mouse weight throughout behavioral tests. *n* = 13-15 per group, **** *P* < 0.0001. Details on statistics provided in **Supp. Table 1**.

### 2.4 Activity Chambers

Mice were placed in individual, closed (but well-ventilated) sound-isolating activity chambers (Med Associates Inc., #ENV-510, 27 × 27 × 20cm). Gross locomotor activity was measured across a 30 minute (min) trial, and these measures are presented (**Fig. 2**): Cumulative Locomotion, Ambulatory Episodes, Ambulatory Time, and Mean Velocity [25].

**Figure 2.**
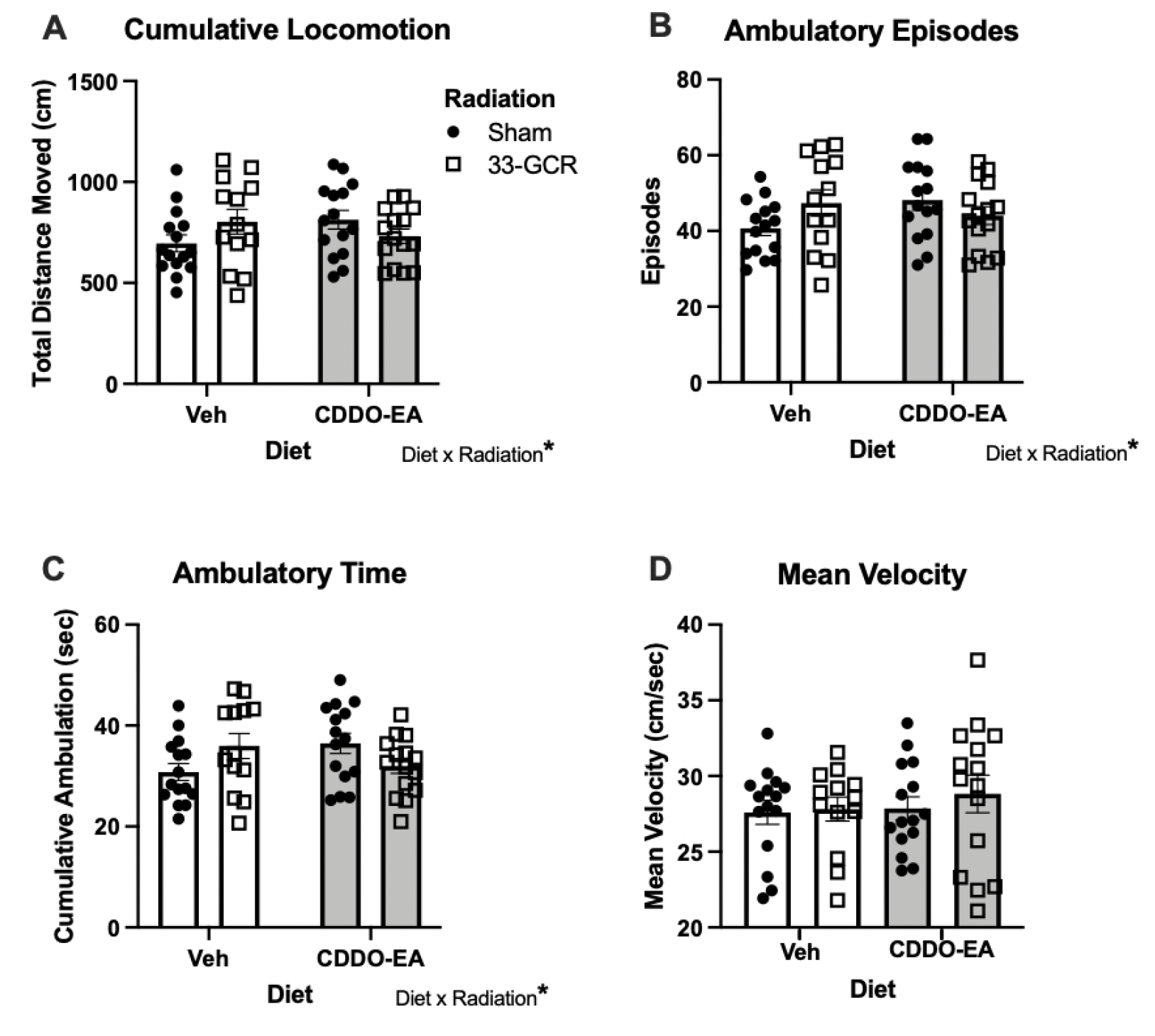
Exposure to 33-GCR and/or CDDO-EA did not change gross locomotor activity measures 4 months post-irradiation. Despite a Diet x Radiation interaction in **A**) Cumulative Locomotion, **B**) Ambulatory Episodes, and **C**) Ambulatory Time (**A-C**), mice in all groups were similar in these measures. **D**) Mean Velocity was also similar among the four groups of mice. *n* = 13-15 per group,* *P* < 0.05. Details on statistics provided in **Supp. Table 1**.

### 2.5 Elevated-Plus Maze (EPM)

Rodent exploratory behavior in the EPM (Harvard Apparatus, #760075) was used as an index of anxiety-like behavior. Each mouse was placed on one of the open arms (43 × 33, 6cm [height x arm length x arm width]) pseudorandomly and allowed free exploration for 5-min [26]. Arena zones were defined as open and closed arm perimeters, and a center zone in between the open and closed arms. Measures presented are Open Arm Exploration, Open Arm Entries, Open Arm Exploration Ratio (ER), and Locomotion (total distance moved; **Fig. 3**). The Open Arm ER for each mouse was defined as:

**Figure 3.**
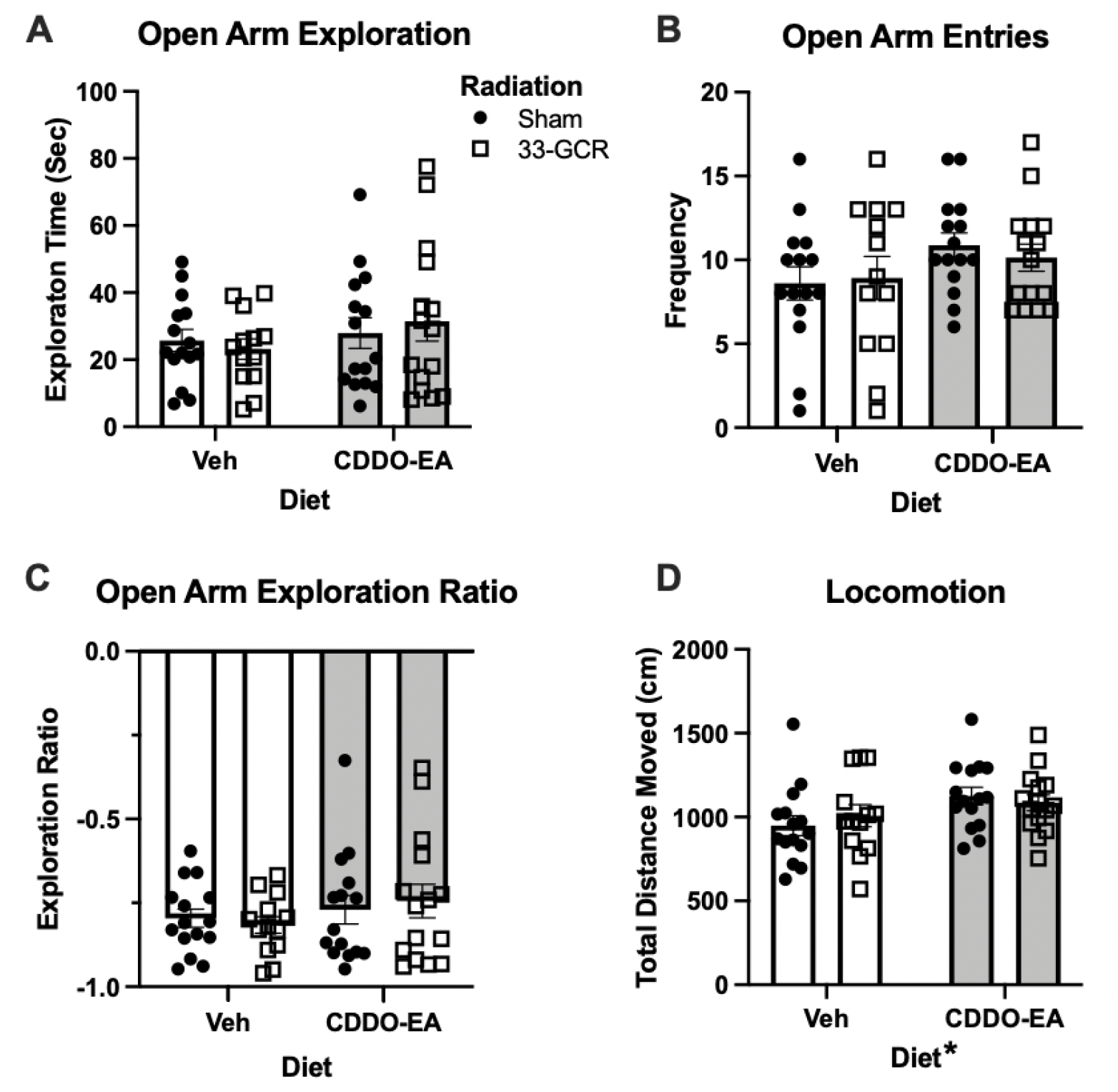
Exposure to 33-GCR and/or CDDO-EA did not change measures in the Elevated Plus Maze 4.5 months post-irradiation. All four groups of mice had similar measures of Open Arm Exploration (Time, **A**), Open Arm Entries (Frequency, **B**), Open Arm Exploration Ratio (**C**). (**D**) In Locomotion (Total Distance Moved), there was a main effect of Diet; mice that received CDDO-EA 4.5 months prior (CDDO-EA/Sham and CDDO-EA/33-GCR mice) moved more than any mice that received Veh (Veh/Sham and Veh/33-GCR). *n* = 13-15 per group, * *P* < 0.05. Details on statistics provided in **Supp. Table 1**.

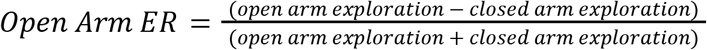

### 2.6 Open Field

Open field exploratory behavior (an index of anxiety-like behavior) was probed by a one-day, 5-min open field paradigm [27]. Mice were placed in the center of a 42 × 42 × 42cm opaque white polycarbonate arena (Nationwide Plastics) for a 5-min trial. General locomotor and exploratory activity was recorded. Behavioral testing was simultaneously conducted on a per cage basis using up to 4 individual adjacent arenas. Mice were placed in the center of the arenas in pseudorandomized orientations. An arena center exploration zone (20 x 20cm) and corner zones (5 × 5cm) were used in analyses. Measures presented are Exploration (time spent in arena center), Thigmotaxis (time spent in corners), Open Field ER, and Locomotion (total distance moved; **Fig. 4**). Open Field ER was defined for each mouse as:

**Figure 4.**
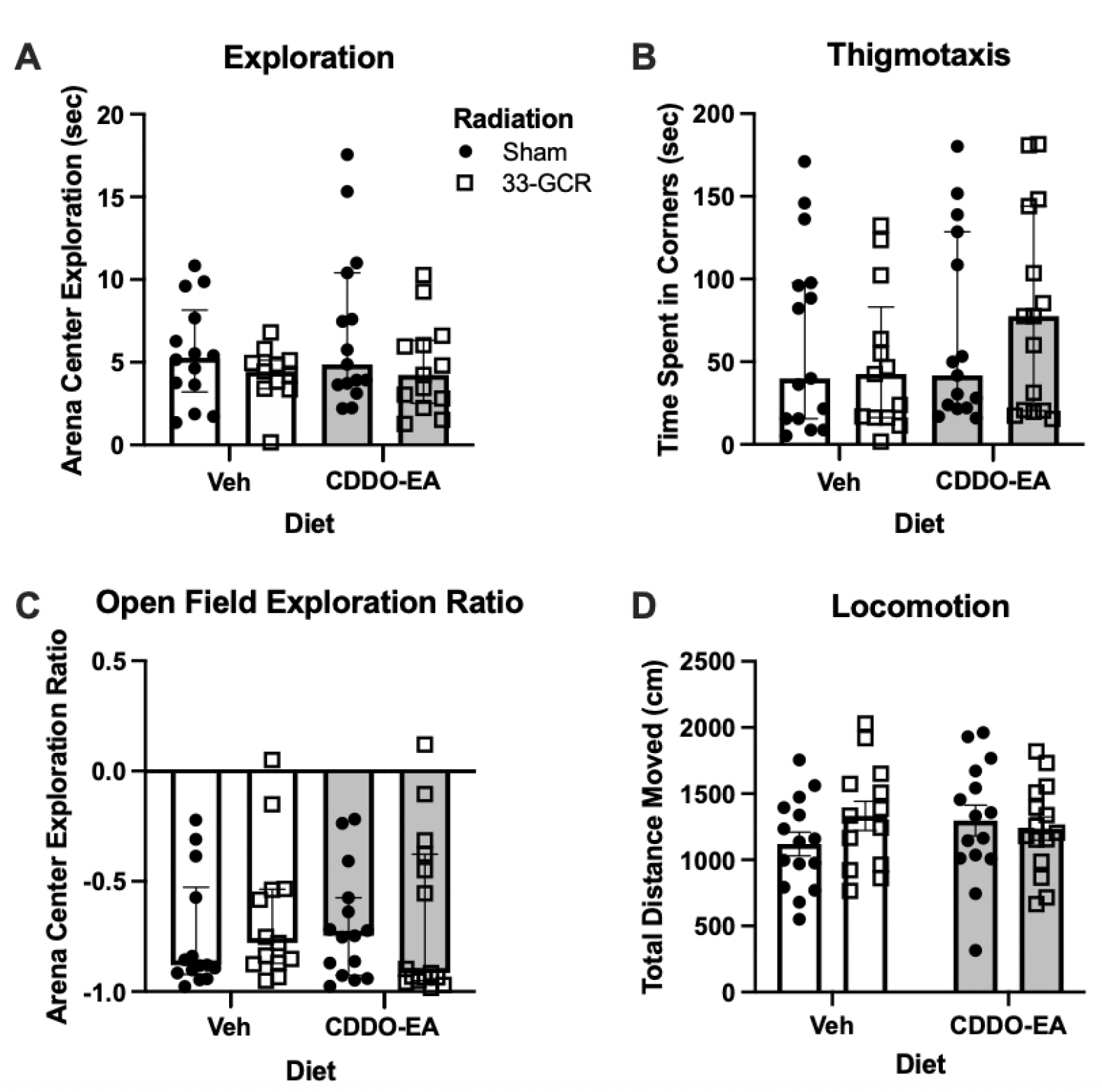
Exposure to 33-GCR and/or CDDO-EA did not change measures in the Open Field 4.5 months post-irradiation. Measures of **A**) Arena Center Exploration, **B**) Arena Corner Exploration, **C**) Exploration Ratio between Arena Center and Corner were statistically similar across experimental groups. **D**) No difference in Total Distance Moved was observed during open field testing. *n* = 13-15 per group. Details on statistics provided in **Supp. Table 1**.

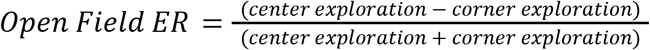

### 2.7 Three-Chamber Social Interaction (3-CSI)

Habituation to a novel arena, sociability, and preference for social novelty were probed by three subsequent trials in the Crawley 3-CSI test [28]. The arena and holders were constructed in the lab out of opaque polycarbonate and acrylic glue. Each chamber measured 40 length × 33 width × 20cm height, with 8cm wide openings to allow movement from the center to the lateral chambers. Each columnar holder was 7cm in diameter, closed on the top and bottom, with 16 evenly-spaced bars (3mm diameter × 16cm height) surrounding the center point. Each test mouse was placed in the center chamber of a 3-chamber arena and allowed free exploration for 10min in each of three consecutive trials (Habituation, Sociability, and Preference for Social Novelty) run on the same day. Inter-trial interval was 1min. Unfamiliar or “novel” mice used for Sociability (Novel Mouse 1) and Preference for Social Novelty (Novel Mouse 1, Novel Mouse 2) trials were strain-, age-, sex-matched, had never had previous contact with test mice, and were group-housed in a separate (but identical) closed-airflow cage system as test mice. For each test subject, the specific novel mice and their locations were pseudo-randomized. For Trial 1 (Habituation) test mice were placed in the empty arena and allowed to explore the three chambers (Left, Center, and Right). Trial 2 (Sociability) involved placing an unfamiliar conspecific stranger mouse (Novel Mouse 1) inside one of the two counterbalanced “holders” in a lateral chamber; the other lateral chamber housed an empty holder, termed here a “Novel Object”. Trial 3 (Preference for Social Novelty) involved leaving the now-familiar Novel Mouse 1 in the same chamber and holder, but another conspecific stranger (Novel Mouse 2) was added to the holder in the opposite lateral chamber. Chamber zones were established by the maze walls, and interaction zones (or “sniff zones”) were defined as 3cm areas surrounding the holders (which in Preference for Social Novelty trial contained the now familiar Novel Mouse 1 or Novel Mouse 2). For Sociability and Preference for Social Novelty trials, the holders were not in direct view from the opposite lateral chamber. Due to the potential confound of contamination of olfactory cues in this test, handler gloves were changed in between each test subject and cage, and the mouse holders and arena were thoroughly cleaned in between subjects and cages. Measures collected were Chamber Exploration Time (for Habituation and Sociability trials) and time spent in sniff zones (for Preference for Social Novelty trial; **Fig. 5**).

**Figure 5.**
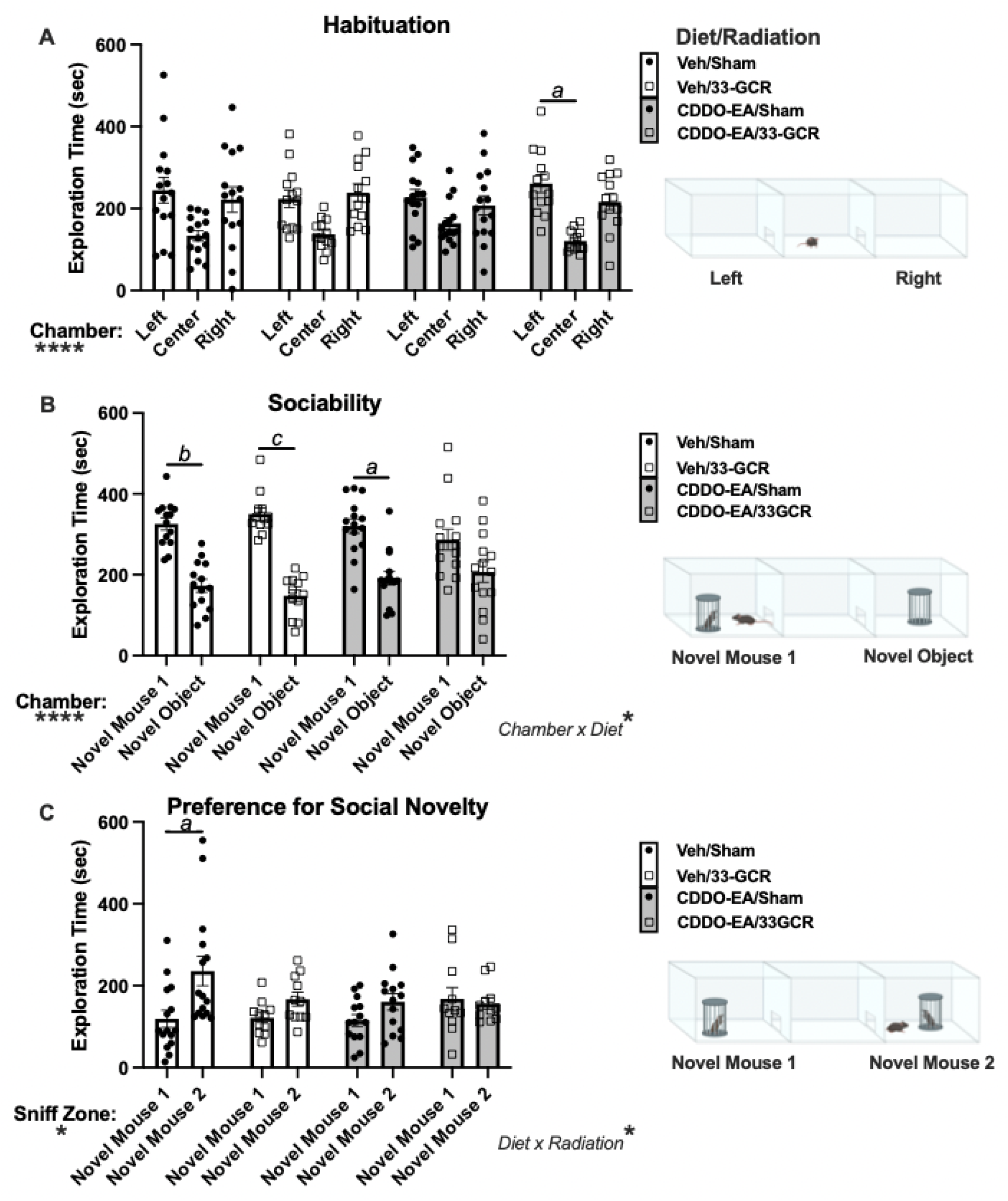
In the 3-chamber social interaction (3-CSI) test, social behavior was altered by 33-GCR and CDDO-EA. Data and schematics depicting experimental conditions for each of the three trials of 3-CSI: **A**) Habituation, **B**) Sociability, and **C**) Preference for Social Novelty. **A**) Habituation to the arena (Left, Center, Right chambers). Of the four groups, only CDDO-EA/33-GCR mice spent more time in one of the lateral chambers (Left) vs. center chamber. However, no group spent more time in one of the lateral chambers vs. the other. **B**) In the Sociability trial, mice in three treatment groups (Veh/Sham, Veh/33-GCR, CDDO-EA/Sham) spent more time in the lateral chamber containing Novel Mouse 1 vs. the Novel Object (an empty holder). CDDO-EA/33-GCR mice spent similar time in both lateral chambers, suggesting a sociability deficit. C) In the Preference for Social Novelty trial, Veh/Sham mice spent more time in the lateral chamber containing Novel Mouse 2 vs. the now-familiar Novel Mouse 1). Veh/33-GCR mice were borderline in spending more time near Novel Mouse 2 vs. Novel Mouse 1. Both CDDO-EA groups (CDDO-EA/Sham, CDDO-EA/33-GCR mice) spent equal time near Novel Mouse 2 and Novel Mouse 1. *n* = 11-15 per group. Main effects: * *P* < 0.05, ** *P* < 0.01, *** *P* < 0.001, **** *P* < 0.0001. Multiple Comparisons: *a P* < 0.05, *b P* < 0.01, *c P* < 0.001. Details on statistics provided in **Supp. Table 1**.

### 2.8 Novel Object Recognition (NOR)

Mice were reintroduced to the open field arena for habituation trials across 2 days [29,30] for 10min free exploration of the empty arena. On day three, mice were introduced to two identical objects (50mL plastic centrifuge tube filled with blue nitrile gloves and water) and allowed free exploration for 10 min. On day four, one object was swapped for a novel object: a 200mL polycarbonate cell culture flask filled with blue aquarium pebbles. For all days, a given mouse was placed in the center of the same arena yet facing a random direction, and on the object recognition test day (day 4) the novel object location was counterbalanced across subjects. Object exploration zones were defined as a ~3 cm perimeter around the external edge of each object. Measures presented are Object Recognition (exploration time near familiar vs. novel object), and Object Discrimination (ratio, **Fig. 6**). Object discrimination ratio (DR) for each mouse was calculated as:

**Figure 6.**
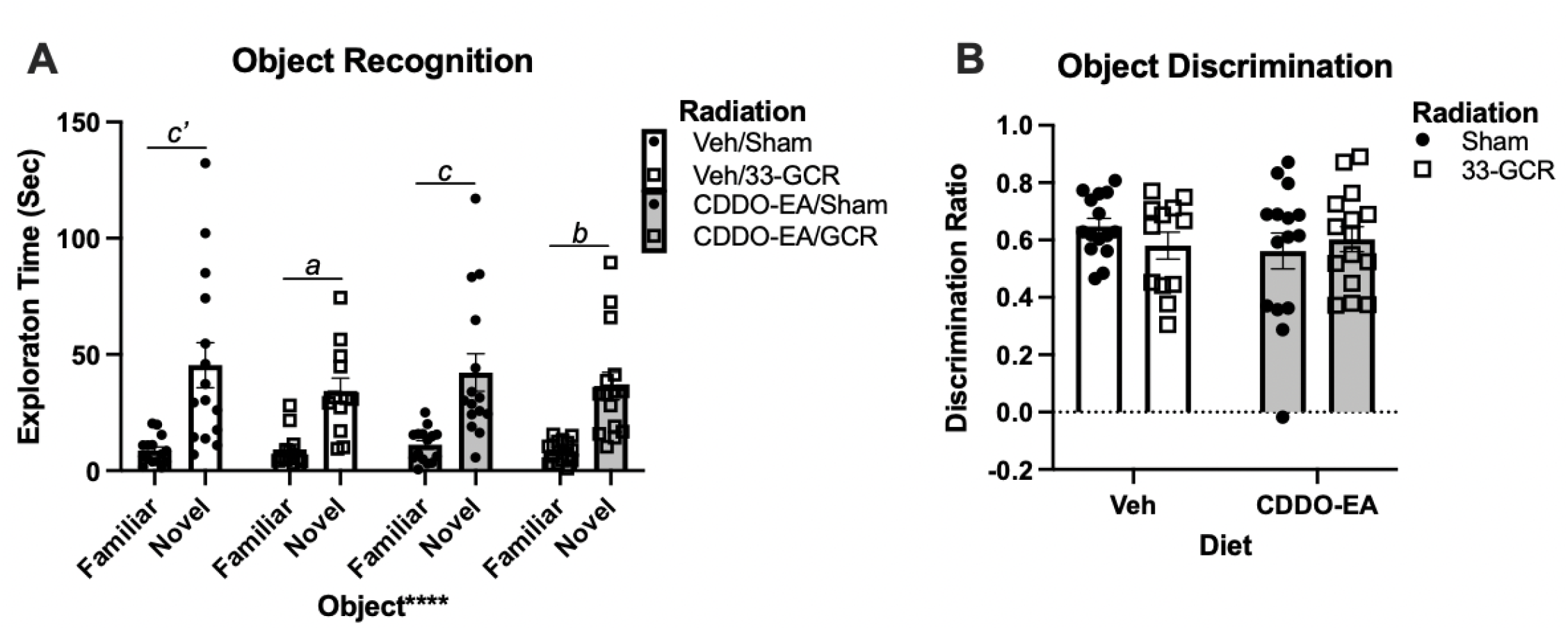
In the Novel Object Recognition task, mice exposed to 33-GCR and/or CDDO-EA performed similarly to control mice. **A**) During the test session, mice in all treatment groups spent more time near the novel vs. familiar object, indicating intact novel object recognition. **B**) All treatment groups also had positive Discrimination Ratios, indicating intact object memory. *n* = 12-15 per group. Main effects: **** *P* < 0.0001. Post-hoc comparisons: *a P* < 0.05, *b P* < 0.01, *c P* < 0.001, *c’ P* < 0.0001. Details on statistics provided in **Supp. Table 1**.

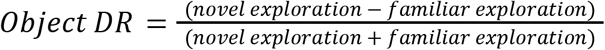

### 2.9 Statistical Analyses and Schematics

Statistical analyses were performed using Prism 9 (Graphpad) unless otherwise specified. Detailed statistical information is provided in **Supp. Table 1**. For all data except for that used for Principal Component Analysis (PCA), outlier testing was first performed via the robust regression outlier method. Next, normality of data distribution was assessed by the Shapiro-Wilk test. Normally-distributed measures were further tested via parametric analyses including 2-way ANOVA (main effects: Radiation, Diet) or 3-way ANOVA (main effects: Radiation, Diet, and Chamber or Sniff Zone) when applicable. Non-normally distributed measures were assessed by Kruskal-Wallis (a non-parametric alternative to a one-way ANOVA) and include a main effect of Treatment rather than Drug and Radiation. H distribution is provided for these nonparametric analyses. For the 3-CSI test, Habituation data were assessed by Repeated Measures (RM) ANOVA, with Chamber (Left, Center, Right) as the RM, and Diet and Radiation as main variables with Diet x Radiation interactions. To assess if time in the center chamber correlated with time in lateral chambers or sniff zone in the Sociability and Preference for Social Novelty trials, Pearson Correlation coefficients were calculated between the time spent exploring the center chamber and the difference (delta) between time spent in the chamber containing Novel Mouse 1 vs Novel Object (Sociability trial) or sniff zones near Novel Mouse 1 vs Novel Mouse 2 (Preference for Social Novelty trial). To determine an effect of Diet or Radiation or possible interactions in each of the three trials of the assay, we applied a 3-way ANOVA with interaction on the difference of time spent exploring the left vs right chambers as the outcome variable. If a significant difference was observed in multiple comparisons in the three trials of the 3-CSI or NOR tests, effect sizes were then calculated by performing post-hoc paired t-tests comparing the time spent in each of the two lateral chambers (Habituation, Sociability), sniff zones (Preference for Social Novelty), or object exploration zones (NOR). Post-hoc multiple pairwise comparisons were performed with Tukey or Dunn’s corrections for parametric and nonparametric measures, respectively. Effect sizes were expressed as percent variance in 2- and 3-way ANOVA analyses, where applicable, and Cohen’s d (provided as *d*) for post-hoc paired t-tests. For mouse survival/attrition rate, the effect of Diet (Veh vs. CDDO-EA) and Radiation (Sham vs. 33-GCR) were first assessed by Cox Proportional Hazards analyses were performed using a custom python script (github.com/EischLab/19AMalesHazardRatios) with the statistics package lifelines [31]. Data are presented as hazard ratio (HR; range 0 to 2), coefficient (*b*_*variable1*_, etc.; range −1 to 1), and probability (*P*). Mouse survival among the four groups was then assessed by the Mantel-Cox Log-rank test. The outcome of these analyses were χ^2^ and *P*. PCA and Pearson’s correlation matrices were used for multivariable behavioral analyses (**Fig. 7**). Principal components were based on parallel analyses, with loadings as behavioral measures for each sample, and with attention to percent variability per leading principal component. Data are expressed as mean +/− SEM when parametric tests were used, and median +/− interquartile range in nonparametric measures. For main effects and interactions, * *P* < 0.05, ** *P* < 0.01, *** *P* < 0.001, and **** *P* < 0.0001. For post-hoc and multiple comparisons, *a P* < 0.05, *b P* < 0.01, *c P* < 0.001, and *c’ P* < 0.0001. α = 0.05. Significant *P* values are italicized in the main text. Schematics were generated with biorender and pasted into Graphpad Prism. Complete details of all analyses (except PCA) are provided in **Supp. Table 1**.

**Figure 7.**
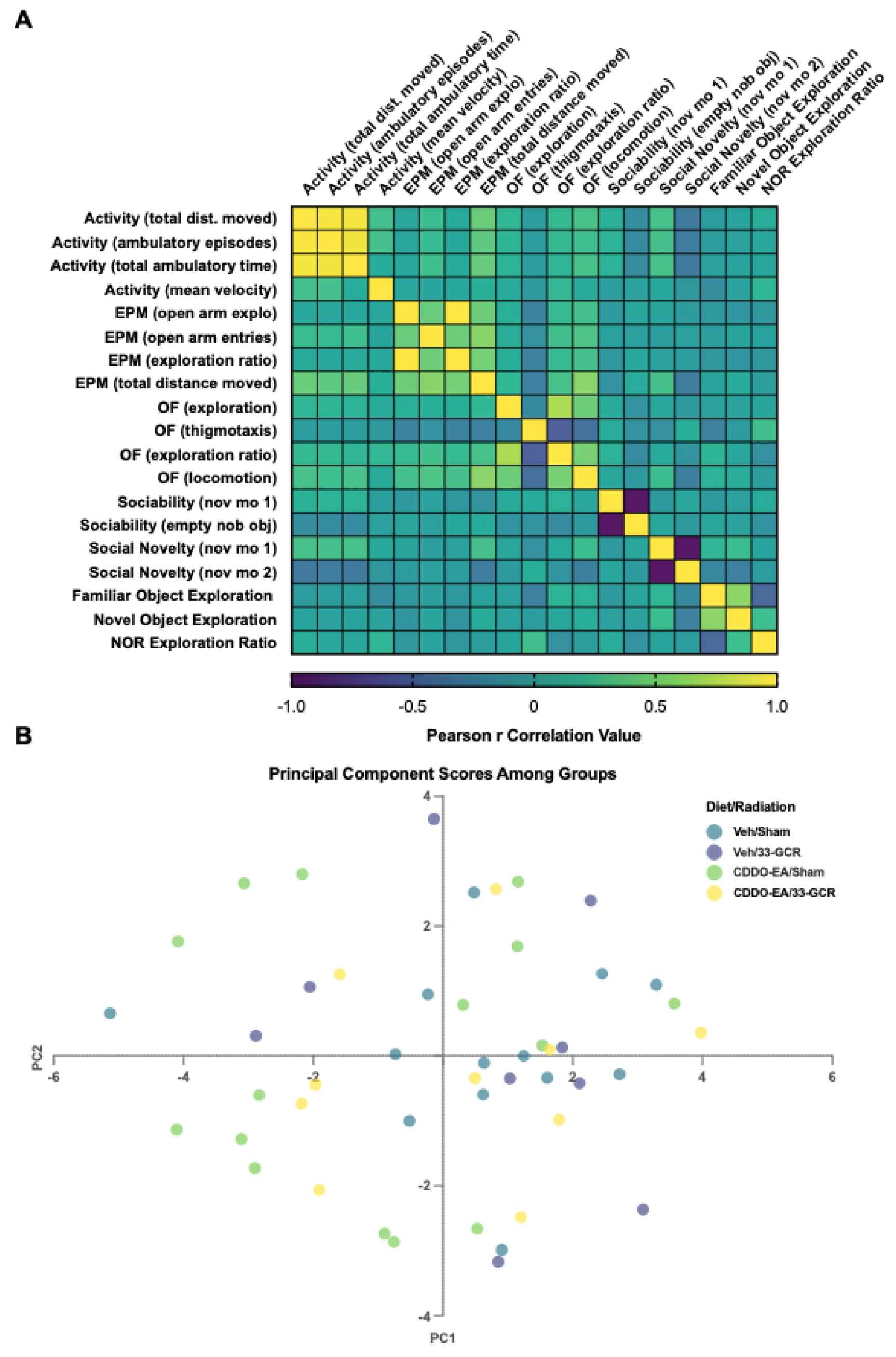
Relationship between behavioral performances between behavioral tests, and variance-based dimensionality reduction among cohorts. **A**) A correlation matrix suggests strong correlations between Activity measures, and moderate correlations between Anxiety, and Exploratory measures. **B**) Principal Component (PC) Analysis of behavioral measures between treatments. A lack of treatment-dependent clusters suggests no gross variance-based behavioral relationships due to specific CDDO-EA or 33-GCR treatments. *n* = 11-15; PC1 = 25.9%, PC2 = 15.56%.

## 3. RESULTS

### 3.1 Study attrition

Rodent use at NRSL is typically accompanied by some level of attrition, likely due to the stress of shipping from a commercial vendor or home institution to NSRL [9,32]. **Supp. Fig. 1** shows the “survival” — in this case, attrition or exclusion — of mice in the present study over the course of the experiment (*n* = 22-24 per group at the start of the study). Mice were primarily removed from the study (and therefore counted as a loss) when they were singly-housed due to within-cage aggression and veterinarian recommendations. Survival data indicate a non-significant hazard of both Diet and Radiation (**Supp. Table 1**). Specifically, there is a non-significant increase in mortality due to CDDO-EA (*b*_CDDO-EA_ = 0.02, *P* = 0.94) or 33-GCR (*b*_33-GCR_ = 0.12; *P* = 0.71). Therefore, there are negligible effects of CDDO-EA and 33-GCR on attrition. When comparing all four groups together, there is no difference among mouse survival curves (χ^2^ = 2.03; *P* = 0.57; **Supp. Table 1**). Thus even though the survival of Veh/33-GCR mice visually appears lower than all other groups, all four groups have a similar attrition rate.

### 3.2 Weights of behaviorally-tested mice

Mouse weights presented in **Fig. 1B** and data in all subsequent figures are from behaviorally-tested mice which, when examined alone, have a flat survival curve (*n* = 13-15 per group; **Supp. Fig. 1**). Since mice typically gain weight throughout adulthood, a predictable main effect of Time is seen in these behaviorally-tested mice (F(16, 448) = 213.5; *P < 0.0001*). However, there is no main effect of Diet (F(1, 406) = 2.208; *P* = 0.14) or Radiation (F(1, 28) = 0.9108; *P* = 0.35; **Fig. 1B**), and no interactions among Time, Diet, or Radiation (**Supp. Table 1**).

### 3.3 Activity Chambers

Four months post-irradiation (**Fig. 1**), mice were tested for gross locomotor activity in fully-enclosed activity chambers (**Fig. 2**). For Cumulative Locomotion (**Fig. 2A**), there is no main effect of Diet (F(1, 54) = 0.2471; *P* = 0.62) or Radiation (F(1, 54) = 0.07420; *P* = 0.79), but there is a Diet x Radiation interaction (F(1, 54) = 4.160; *P = 0.0463*). Post-hoc multiple comparisons reveal no differences beyond chance in Cumulative Locomotion among the four groups (**Supp. Table 1**). The same was true for Ambulatory Episodes (**Fig. 2B**): no main effects of Diet (F(1, 54) = 0.6358; P = 0.43) or Radiation (F(1, 54) = 0.2526; *P* = 0.62), but a Diet x Radiation interaction (F(1, 54) = 4.268; *P = 0.044)* with no multiple comparison-based differences (**Supp. Table 1**).This was also true for Cumulative Ambulation (**Fig. 2C**): no main effect of Diet (F(1, 54) = 0.2024; *P* = 0.66) or Radiation (F(1, 54) = 0.03055; *P* = 0.86), a Diet x Radiation interaction (F(1, 54) = 6.410; *P = 0.01*), and no multiple comparison-based differences (**Supp. Table 1**). The significant interaction between Diet and Radiation suggests that although the effect of Diet is not significant overall, it differs between radiation groups. Mean Velocity — the one non-parametric analysis performed on activity chamber data — was also not different among groups (**Fig. 2D**; H = 1.381; *P* = 0.71; **Supp. Table 1**).

### 3.4 Elevated Plus Maze (EPM)

Mice are naturally reluctant to leave an enclosed area for an open area, and thus spend more time in the closed vs. open arms in the EPM. However, due to endogenous exploratory drive, they will occasionally overcome their putative exposure anxiety and venture into the open arms, providing a quantifiable measure of anxiety-like behavior (**Fig. 3**). In open arm exploration time, there are no main effects of Diet (F(1, 54) = 1.398; *P* = 0.24) or Radiation (F(1, 54) = 0.01066; *P* = 0.92; **Fig. 3A, Supp. Table 1**). Similarly, in open arm entries (H = 2.675; *P* = 0.45; Fig. 3B, **Supp. Table 1**) and Open Arm ER (H = 0.0412; *P* = 0.94; **Fig. 3C, Supp. Table 1**) are not different among groups. Since anxiety-like activity often manifests in freezing behavior, cumulative locomotion of mice was also recorded throughout the entire EPM test period (**Fig. 3D**). In cumulative locomotion, there is a main effect of Diet (F(1, 54) = 5.134; *P = 0.028*) but not of Radiation (F(1, 54) = 0.03087; *P* = 0.86) and no Diet x Radiation interaction (F(1, 54) = 0.7894; P = 0.37). Mice given CDDO-EA at the time of either 33-GCR or Sham irradiation therefore move more in the EPM vs. mice given Vehicle; however, the size of this effect is relatively small. Multiple comparisons reveal no difference beyond chance in mean total distance moved among the four cohorts (**Supp. Table 1**).

### 3.5 Open Field

Open field behavior was tested next (**Figs. 1, 4**). The open field arenas are distinct from the activity chambers (Section 3.3) as they have no “roof”, and thus promote anxiety-like behavior in a prey species such as mice. Similar to the EPM, mice are reluctant to explore the open portion of the open field and will spend a larger proportion of time in the corners, once more providing a basis for quantifying anxiety-like behavior. There is no evident variation in arena center exploration among groups (H = 1.961; *P* = 0.5805; **Fig. 4A, Supp. Table 1**) or of time spent in the corner of arenas (thigmotaxis; H = 2.122; *P* = 0.5475; **Fig. 4B, Supp. Table 1**). Exploration ratios between time spent in the center vs. corners show no main effect of Treatment (H =1.823; *P* = 0.6100; **Fig. 4C, Supp. Table 1**). In line with measures of exploratory or anxiety-like behavior, open field locomotion analysis shows no main effect of Diet (F(1, 54) = 0.1548; *P* = 0.6955) or Radiation (F(1, 54) = 0.5643; *P* = 0.4558), or Diet x Radiation interaction (F(1, 54) = 1.818; *P* = 0.1832; **Fig. 4D, Supp. Table 1**).

### 3.6 Three-Chamber Social Interaction

Three-CSI consists of three sequential trials — Habituation, Sociability, and Preference for Social Novelty — tested on a single day, with slight difference in chamber set-up in each trial (**Fig. 5**).

#### 3.6.1 Three-Chamber Social Interaction: Habituation

In the Habituation trial, time spent in each of the three empty chambers was measured to assess possible chamber or testing environment bias (**Fig. 5A**). During Habituation, mice may spend more time in the more enclosed lateral (left and right) chambers vs. the center chamber. Time spent in the left vs. right chambers during Habituation is a critical metric to consider, as unequal time in one of these lateral chambers at “baseline” can influence behavior in the subsequent two trials (Sociability and Preference for Social Novelty). 3-way ANOVA on 3-CSI Habituation data suggests a main effect of Chamber (F(2, 104) = 15.92; *P* < 0.0001), but no effect of Diet (F(1, 52) = 1.465; *P =* 0.2361) or Radiation (F(1, 52) = 0.1193; *P* = 0.7312) and no interaction of any sort (**Supp. Table 1**). Post-hoc multiple comparison within-group reveals no significant difference of Chamber in most groups. The exception was the CDDO-EA/33-GCR group which explored the left chamber for a greater duration than the center chamber (*P* < *0.05*; **Fig. 5A**). However, as no group spent more time in the left vs. right chamber, these results overall suggest normal habituation to the three-chamber arena.

#### 3.6.2 Three-Chamber Social Interaction: Sociability Trial

In the Sociability 3-CSI trial, mice are tested for their natural “sociability” to spend more time near Novel Mouse 1 in a holder in one of the lateral chambers vs. an empty holder (considered a Novel Object) in the opposite lateral chamber (**Fig. 5B**). Importantly, lateral chambers were pseudorandomized for the placement of the Novel Object. We first tested within each of the four groups for correlations between the Center Chamber Exploration Time and the difference of time (delta) spent exploring Novel Mouse 1 vs. Novel Object chamber. No significant correlations are observed within each group, suggesting no center chamber exploration bias in lateral chamber exploration (**Supp. Table 1**; raw data **Supp. Fig. 2**). We next tested for the influence of Diet, Radiation, and Chamber (Left, Right, and Center) on chamber exploration time via 3-way ANOVA. There is a main effect of Chamber (F(1, 51) = 55.52; *P* < 0.0001), but no effect of Diet (F(1, 51) = 0.3034; *P =* 0.2361) or Radiation (F(1, 51) = 1.887; *P* = 0.1756; **Supp. Table 1**). Of all interactions, there is only one: a Diet x Radiation (F(1, 51) = 4.205; *P* < 0.05; **Supp. Table 1**). Post-hoc multiple comparison suggests that Veh/Sham, Veh/33-GCR, and CDDO-EA/Veh mice spend more time exploring Novel Mouse 1 vs. the Novel Object, with medium to large effect sizes (*P < 0.01*, *P < 0.0001*, *P < 0.05*, respectively; **Supp. Table 1**). However, CDDO-EA/33-GCR mice show no difference in time spent exploring Novel Mouse 1 vs. the Novel Object (*P* = 0.3502; **Supp. Table 1**). These results indicate that, unlike all other groups, CDDO-EA/33-GCR mice display sociability deficits.

#### 3.6.3 Three-Chamber Social Interaction: Preference for Social Novelty Trial

Finally, mice were tested in the 3-CSI Preference for Social Novelty trial (**Fig. 5C**). This trial was originally used to measure olfactory detection of social odors, as mice typically will spend more time near an enclosed novel vs. familiar mouse [33]. However, as this trial also involves social discrimination, it has been widely used to assay social memory [34–36]. Similar to the Sociability trial, here we tested for center chamber exploration bias by testing for correlations between the time spent exploring the center chamber and the difference of time spent in the sniff zone of an unfamiliar Novel Mouse 2 and the sniff zone of the now-familiar Novel Mouse 1. The difference in sniff zones is interpreted as an index of preference for social novelty, where unimpaired mice are expected to spend more time in the sniff zone of Novel Mouse 2 vs. Novel Mouse 1. Correlation analysis did not detect any significant correlations between Center Chamber Exploration time and time spent in either lateral chamber sniff zone (**Supp. Table 1**; raw data in **Supp. Fig. 2**). Three-way ANOVA reveals a main effect of Sniff Zone (F(1, 47) = 6.595; *P* < 0.05) and one interaction: Diet x Radiation (F(1, 47) = 4.719; *P* < 0.05; **Fig. 5C**). Post-hoc multiple comparison shows Veh/Sham mice spend more time in the Sniff Zone of Novel Mouse 2 vs. Novel Mouse 1 (*P* < 0.05; *d* = 0.2665; **Supp. Table 1**), suggesting normal preference for social novelty. However, none of the remaining groups spend more time in the Sniff Zone of Novel Mouse 2 vs. Novel Mouse 1 (Veh/33-GCR, CDDO-EA/Sham, and CDDO-EA/33-GCR all *P* > 0.05; **Fig. 5C**). Overall, these data show a Diet and Radiation-dependent deficit in preference for social novelty, with 33-GCR and CDDO-EA each on their own leading to less sniff zone time near Novel Mouse 2. Also, the data from CDDO-EA/33-GCR mice show this administration of CDDO-EA does not prevent the 33-GCR-induced blunting of preference for social novelty.

### 3.7 Novel Object Recognition

NOR is another task that takes advantage of a mouse’s exploratory drive to investigate unfamiliar over familiar objects. When analyzing the time mice spent exploring a 3cm margin around each object during the novel object testing trial (**Fig. 6A**), there is a main effect of Object (F(1, 53) = 78.75; *P < 0.0001*), but not of Diet (F(1, 53) = 0.0013; *P* = 0.97) or Radiation (F(1, 53) = 1.149; *P* = 0.23) and no interactions evident (**Supp. Table 1**). Multiple comparisons suggest all four groups explore the novel object more than the familiar object (Veh/Sham: *P < 0.0001*; Veh/33-GCR: *P = 0.023;* CDDO-EA/Sham: *P = 0.0005*; CDDO-EA/33-GCR: *P = 0.002*). To further assess the influence of treatment on object recognition, discrimination ratios were plotted (**Fig. 6B**). All object discrimination ratio of all groups are positive, and there are no main effects of Diet (F(1, 53) = 0.4505; *P* = 0.51) or Radiation F(1, 53) = 0.0754; *P* = 0.71), and no Diet x Radiation interaction, suggesting intact and similar object discrimination across groups (**Fig. 6B**).

### 3.8 Multivariable Behavioral Analyses

PCA determines possible linear combinations of behavioral variables that account for the most variance by reducing the dimensionality of a multivariable dataset. This method allows for the visualization of related behaviors in a treatment-based manner. Individual samples are also displayed by reduced dimensionality of all behavioral values, allowing visualization of related individuals by Diet and Radiation such that clusters emerge in similarly-performing individuals in a manner that does not factor in treatment in the calculation of plot coordinates. The total number of behaviorally-tested mice included in the analyses after outliers exclusions is 47, and the number of behavioral variables is 19. PC1 accounts for a relatively low 25.9% of the total variance, followed by 15.6% in PC2. A Loadings Plot reveals several relationships between behavioral measures (**Supp. Fig. 3**). Anxiety-like and activity measures generally align in a cluster along the PC1 axis (PC1 = −0.45 to −0.85) and appear to be somewhat separate from object and novel mouse exploratory measures (PC1 = −0.2 to 1.0), with open field thigmotaxis and novel mouse 2 exploration isolated. Interestingly, however, PC2 clearly separates anxiety-like measures from activity measures (PC2 = 0 to −0.35, and 0.5 to 0.65, respectively). PC Loadings reveal several close variance relationships. For example, EPM exploration ratio is highly predictive of open field ER, suggesting the variances in height-dependent and open field exposure anxiety-like behavior are highly similar. An unexpected pattern that emerges is that the variance in open field exploration is tightly related to that of the time spent with novel mouse 1 in the preference for social novelty trial of the 3-CSI test, though the two measures do not appear to correlate (**Fig. 7A, Supp. Fig 3**). Indeed, anxiety-like measures broadly show a moderate correlation, whereas exploratory behaviors do not (**Fig. 7A**). PC scores for individual mice reveal no treatment-based clustering, suggesting low general behavioral predictability due to treatment (**Fig. 7B**).

## 4. DISCUSSION

To investigate the potential effects of a ground-based 33-beam GCR analog exposure on the CNS of mature male mice, we compared their behavioral performance on a variety of tasks starting 4 mo after exposure to an acute, 75cGy, Mars mission-relevant dose of whole-body 33-GCR or Sham irradiation. We additionally assessed whether a transient, five-day diet of CDDO-EA (400mg/Kg) or Vehicle given before, during, and immediately after irradiation prevented or attenuated behavioral changes associated with radiation. Our most striking findings relate to the social domain: sociability in 3-CSI was compromised in CDDO-EA/33-GCR mice and preference for social novelty was blunted in Veh/33-GCR and CDDO-EA/Sham mice. Also of note is that the potential countermeasure, CDDO-EA, given in chow before, during, and immediately after irradiation did not attenuate the 33-GCR-induced blunting of preference for social novelty. Surprisingly, 33-GCR had no effect on a hippocampal-dependent task, NOR, and did not increase anxiety-like behaviors (assessed in open field and EPM); CDDO-EA also did not influence these measures. These behavioral results — which to our knowledge are the first from rodents exposed to the complex 33-ion GCR analog now available at NASA’s Space Radiation Laboratory — are discussed in the context of the much wider literature on the behavioral impact of single-, multiple-, and sequential-particle exposures on rodents and for their implications for deep space travel.

Whole-body exposure to Mars-relevant doses of charged-particle radiation often leads to deficits in rodent cognition and altered behavior [c.f. 4,5–7,37]. Among the most commonly researched and reported effects is that charged-particle radiation decreases rodent NOR, an ability linked to hippocampal integrity [38]. The numerous reports of charged-particle irradiation-induced NOR deficits contribute to the long-standing belief that the hippocampus is a particularly radiosensitivity brain region. NOR deficits are even evident relatively long after irradiation. For example, whole-body exposure to sequential mixed beams (50cGy protons, 1hr break, 10cGy ^16^O) decreased NOR in male mice 3 and 9 mo post-irradiation, and decreased object memory at the 9 mo time point [30,39]. Disruptions in other hippocampal-dependent functions are also reported in male mice after exposure to lower energy proton irradiation as well [40,41]. The preponderance of published deficits in hippocampal-dependent tasks after charged-particle irradiation make it all the more surprising that here we report no 33-GCR-induced changes in NOR 6 mo post-irradiation. Additional evidence that 33-GCR does not grossly impact hippocampal function comes from the similar levels of exploration and locomotion we report in 33-GCR mice relative to other groups; if 33-GCR induced frank hippocampal lesions, 33-GCR mice would have been hyperactive [42–45]. We also report here that 33-GCR does not change anxiety-like behavior (open field, EPM), which is consistent with some simple and sequential mixed beam charged particle studies [30] but not others [11,46]. As with many dependent variables, the discrepancies in open field results among labs could be due to a host of experimental differences, from radiation differences to husbandry practices to open field size and test duration. Efforts to standardize testing - or to normalize data collection and analysis, perhaps via optical imaging and machine learning - may allow more direct comparison of results from different laboratories. While our 33-GCR mice were similar to control mice in the hippocampal-dependent NOR task and anxiety-like behaviors, 33-GCR mice did show a blunted preference for social novelty 5.5mo post-irradiation. Due to the extensive social challenges associated with deep spaceflight, this finding with 33-GCR is notable in that it is the first to suggest a social domain deficit in rodents after exposure to a Mars-relevant dose of a highly-complex mixed beam paradigm. Our work with 33-GCR is in line with prior work with single or sequential mixed beams. For example, in male rats or male and female mice, single or mixed beam (^1^H, ^16^O, and ^28^Si) whole-body 10-50cGy exposure decreases preference for social novelty 1 to 9 mo post-irradiation [9,47,48]. One potential caveat is that the social novelty trial of the 3-CSI is known for its high inherent variability; perhaps the lack of difference in many of the groups means this trial of the 3-CSI is not working optimally. This concern is somewhat ameliorated by two things: 1) Our analysis includes effect sizes, providing insight into the actual magnitude of any detected differences; this is in contrast to *P* which does not reflect magnitude of difference [49–53]. 2) Notably, the Veh/Sham group spent more time exploring the Novel Mouse 2 vs. 1, and did so with a small-moderate effect size, suggesting the trial is working well (at least for these control mice). Given that the social novelty trial of the 3-CSI was initially designed to test olfaction of social odors [33], future work with 33-GCR ideally will involve testing for olfaction and olfaction discrimination of both social vs. non-social and novel vs. familiar cues in order to address these and related possible confounds.

Why might the 75cGy dose of 33-GCR paradigm used here blunt preference for social novelty in male mice, but not cause the changes in anxiety-like behavior, object recognition memory, or sociability seen in prior studies with similar post-irradiation time points and similar doses (albeit of single- or simple mixed particles) [8]? One reason could be that preference for social novelty is among the first behavioral phenotypes to decline with age in mice [54], and, as previously suggested, charged-particle radiation may accelerate aging phenotypes [55–58]. In this regard, it is interesting to compare the impact of mixed and 33-GCR exposure on social behaviors when tested at different intervals after irradiation. For example, in our present work, mature (6 mo old) male mice exposed to 33-GCR (75cGy) show normal sociability yet worse preference for social novelty when tested 5.5 months later (at ~11.5 mo of age). This is in contrast to other work where younger mature (4-4.2 mo old) male mice exposed to 3-beam GCR (50cGy) have worse sociability and borderline worse preference for social novelty when tested 1.5-2 months later (at 5.5-7 mo of age) [9]. Future work with 33-GCR exposure should closely consider the relationship between radiation and aging pathologies, as has been done in-depth with previous single particle exposures in rats [59]. Another possible reason is that a complex mixed radiation field (as used in the present study) has distinct radiation properties relative to single- or simple mixed-particle exposures. In this regard, it is useful to consider a peculiar aspect of radiation interaction with tissues: the dependence of Linear Energy Transfer (LET) of specific particles on type of DNA damage and damage response. Low-LET ^4^He radiation induces a far lower proportion of clustered lesions when compared to high-LET ^4^He, the damage of which consists almost entirely of clustered lesions as shown by exposure modelling [60,61]. In addition, proton-induced DNA double-strand breaks are also LET-dependent, which correlate with the relative biological effectiveness of the cell [62]. DNA damage response is likewise LET-dependent. When comparing low- to high-LET proton and ^4^He exposures on varying non-neuronal cancerous cells, there are distinct LET-dependent mechanisms of recognition and response to DNA damage [63,64]. With respect to particles of similar LET, double-stranded DNA breaks increase as particle track radius increases, which is dependent on the particle’s kinetic energy [65]. Based on these observations, a complex particle field such as that of 33-GCR — which involves a combination of low- and high-LET radiation and particles of varying Z — is expected to recruit repair machinery primarily associated with the high-LET components of the radiation field. This may help explain why mice exposed to 75cGy of 33-GCR have fewer changes in behavior relative to mice exposed to similar doses of only high- or low-LET radiation [20]. Furthermore, the cell-based studies showing LET-dependent differences in DNA repair used very high doses. It is also possible that there are “upside-down-U” dose-dependent effects of radiation repair where lower doses are not as effective at DNA repair response, as previously suggested [66]. This may explain why a 30cGy dose of 5-GCR decreased performance of mice in several behavioral domains [11] whereas the 75cGy dose of 33-GCR used here only decreased preference for social novelty. Future work using 33-GCR at different doses should elucidate the potential dose- and LET-properties of a complex radiation field on DNA damage and CNS response.

While mechanistic underpinnings of the 33-GCR-induced deficits in preference for social novelty were outside the scope of this study, it is reasonable to consider that the preference for social novelty deficits were due to brain circuit-specific radiosensitivity. Although the exact circuit-dependence for neurotypical sociability remains uncertain, optogenetic and DREADD manipulation studies implicate basal forebrain-ventral tegmental area (VTA), cerebellum-VTA, basolateral amygdala-nucleus accumbens, and medio-dorsal thalamus-medial prefrontal cortex tracts as critical for sociability as assessed in the 3-CSI task [67–70]. These circuits are distinct from those that underpin preference for social novelty in 3-CSI: lateral entorhinal cortex, median raphe nucleus, and hypothalamic supramammillary nucleus, and specifically the hippocampal CA2 [71,72]. Of these distinct circuits, CA2 is of particular note; it is unique among hippocampal subregions in its projection pattern within the hippocampus and to non-hippocampal regions [73]. In fact, disruption of CA2 interferes with preference for social novelty but leaves both sociability and NOR intact [71]. Therefore, it is reasonable to speculate that the 33-GCR findings presented here — worse preference for social novelty but not change in sociability and NOR — may emerge from 33-GCR-induced damage to CA2. This concept is indirectly supported by prior work: 1) 25cGy of ^16^O impaired preference for social novelty, profoundly reduced the length of CA2 pyramidal neuron dendrites, and decreased both dendritic complexity and CA2 mushroom spine density in mature female mice [48], and 2) CA2 receives monosynaptic projections from the dentate gyrus [73], a hippocampal subregion long known to be sensitive to GCR [e.g. 74,75,76]. We advise that future work specifically interrogates the potential effects of 33-GCR on the hippocampal CA2 and its associated regions. If the hippocampal CA2 is actually more radiosensitive than other hippocampal subregions, dysfunction in CA2-linked behaviors could serve as early indicators of 33-GCR-induced CNS impact.

Due to the complexities and costs associated with radiation shielding, pharmacological countermeasures are being considered to mitigate radiation exposures. One promising family of antioxidant compounds is the triterpenoid oleanolic acid derivative CDDO, which can be orally administered. Several CDDO variations exist with differing moieties on carbon 28. One such compound, CDDO-Methyl, has gone through a number of clinical trials and is at the time of preparation of this manuscript in anti-inflammatory therapy Phase 3 trials (NCT03749447) for chronic kidney disease. CDDO is of particular interest for CNS research due to effective pre-clinical therapeutic intervention for mouse models of Huntington’s disease, malaria, and ischemic injury [16–19]. In addition, dietary CDDO-EA (400mg/Kg) is an effective countermeasure against

GCR-induced lung tumors in tumorigenic male mice exposed to 30cGy of a 3-beam mixed field [15]. CDDO-EA was therefore a promising intervention as it offers a multifaceted protective approach by targeting the Nrf-2 pathways, which have protective downstream effects on oxidative stress management, microglial activation, and blood-brain barrier integrity, all of which are adversely affected in the CNS of mice following radiation [77,78]. Contrary to our expectations, CDDO-EA did not protect male mice from 33-GCR-induced deficits in social novelty. However, the mice received CDDO-EA transiently: before, during, and immediately after 33-GCR exposure. This is in contrast to ongoing human clinical trials where volunteers receive daily oral CDDO-Methyl (Bardoxylone). It is possible that CDDO-EA may exert a radioprotective effect if different administration parameters are employed (longer duration, higher dose) or if used under even more physiologically-challenging radiation exposures (higher dose, greater proportion of dose is from high LET). Indeed, prior studies with dietary antioxidants show mice and rats are protected from radiation-induced behavioral impairments, but only when the radiation dose was relatively high (>150cGy) [23,79–81]. Future work is warranted to understand the intricacies of the interactions between 33-GCR exposure, CDDO-EA administration parameters, and behaviors in the social domain.

There are limitations to the current work. The first and most important limitation is that it was performed in male mice, and thus provides no insight into how 33-GCR impacts these behaviors in female mice. This is particularly important limitation since over half the current astronaut class is female and there are sex-dependent differences in CNS sensitivity to charged particle radiation [9,82]. It is also important from a fundamental neurobiological perspective; female and male rodents differ in key developmental and behavioral aspects, in their response to stress, and in both their performance in and strategies for completing many “classical” behavioral paradigms [83–87]. Indeed, given the lower level of within-cage aggression in group-housed conspecific female vs. male mice (see next point), studies on female mice likely will have lower attrition than is reported here and elsewhere in male rodents [9]. The essential future experiments on the sex-dependent impact of space radiation on cognition and behavior are made possible by the recent reiteration of the appropriate study design and the exaggeration of the impact of female hormones [88,89]. A second limitation is that attrition rates in this study (35-55%) are higher than previous reports (5-15%) [9,32]. It is important to note that the attrition is primarily due to within-cage aggression subsequent veterinarian recommendations to individually-house certain mice. No behavioral data from single-housed mice are included in this study given the known behavioral impact of the stress of solo housing [90–92]. In the present study, mice were ordered and shipped directly to NSRL from the breeder. In our past work, mice were first shipped to our home institution and then to NSRL and back, and this resulted in much lower attrition. Further, when stressed, male mice are more likely to exhibit aggressive behaviors, whereas an inverse relationship between plasma corticosterone and aggression occurs in females, suggesting sexually dimorphic responses to stress under normal housing conditions [92,93]. An important confound related to the inclusion of subjects in studies requiring multiple transport — such as in the current study — is that transport induces stress in mice, which may interact with the response to a manipulation (such as irradiation), and that this may manifest in a sex-specific manner. This is yet another reason why the current study should be repeated in female mice; they are far less prone to the within-cage aggression that led us to single-house (and thus exclude from behavioral analysis) many male mice in our study. A third limitation is the fluence used. In the present study, 75 cGy was delivered in 1-2 hrs. As the GCR fluence in space is on the order of 0.01 cGy/day [94], future work would ideally consider the dose-rate effects of 33-GCR on subsequent behavior changes.

In sum, we present to our knowledge the first data on the effects of whole-body 33-GCR on the CNS. We conclude in male mice that 1) 33-GCR is detrimental to preference for social novelty when tested months after exposure, 2) transient, prior administration of CDDO-EA did not block the radiation-induced preference for social novelty deficit, 3) short-term CDDO-EA given months prior can itself decrease preference for social novelty, and 4) prior CDDO-EA given at the time of 33-GCR together result in suppressed sociability. Also, mature male mice exposed to 33-GCR had normal levels of locomotion, sociability, and even – in contrast to our hypothesis – normal levels of anxiety-like behavior and hippocampal-dependent memory. This work opens several new avenues of investigation. For example, our data encourage future studies on the CNS effects of space radiation to closely consider the role of LET in both cellular (DNA damage recognition and repair promotion) and behavioral aspects. Whereas much research has been done the impact of space radiation on cognitive behaviors [8], more research on social behaviors is warranted due to radiation-dependent changes in sociability and preference for social novelty following exposure to charged-particle radiation [9,47,48] as well as the relevance of social behavior to prolonged spaceflight [95]. Future studies should pursue more translationally-relevant paradigms which limit a number of confounds associated with open arena behavioral testing [37]. Taken together, our data underscore the brain remains a relevant area of concern for Mars-relevant spaceflight.

## 5. ACKNOWLEDGEMENTS

This work was supported by NASA grants 80NSSC17K0060 (Eisch), 80NSSC21K0814 (Yun), NNX16AE08G (Shay); by the Translational Research Institute for Space Health through NASA cooperative agreement NNX16AO69A (Kiffer); NIH MH107945 (Eisch and Yun); the University of Pennsylvania McCabe Fund (Yun); the NARSAD Brain & Behavior Research Foundation award for Young Investigators (Yun); the Children’s Hospital of Philadelphia Department of Anesthesiology and Critical Care Development Fund (Eisch); the Douglass Project at Rutgers University (Patel); Upenn Post-Baccalaureate Education Program (PennPrep) through NIH R25GM071745 (PI: K. L. Jordan-Sciutto) and the Biomedical Graduate Studies at the University of Pennsylvania (Guzman and Soler). This research was also supported by NASA Grants NNX07AP84G (co-I Eisch), NNX12AB55G (co-I Eisch), NNX15AE09G (Eisch); NIH grants DA007290, DA023555, DA016765, MH107945 (Eisch) and R15 MH117628 (PI: K. G. Lambert); and a pilot grant from the University of Pennsylvania Perelman School of Medicine Department of Radiation Oncology (co-I Eisch). Thank you to the staff of Brookhaven National Labs and the NASA Space Radiation Laboratory for experiment support and coordination.

## Data Availability Statement

Experiment data will be made available upon reasonable written request.

## SUPPLEMENTARY DATA

**Supplementary Table 1.**
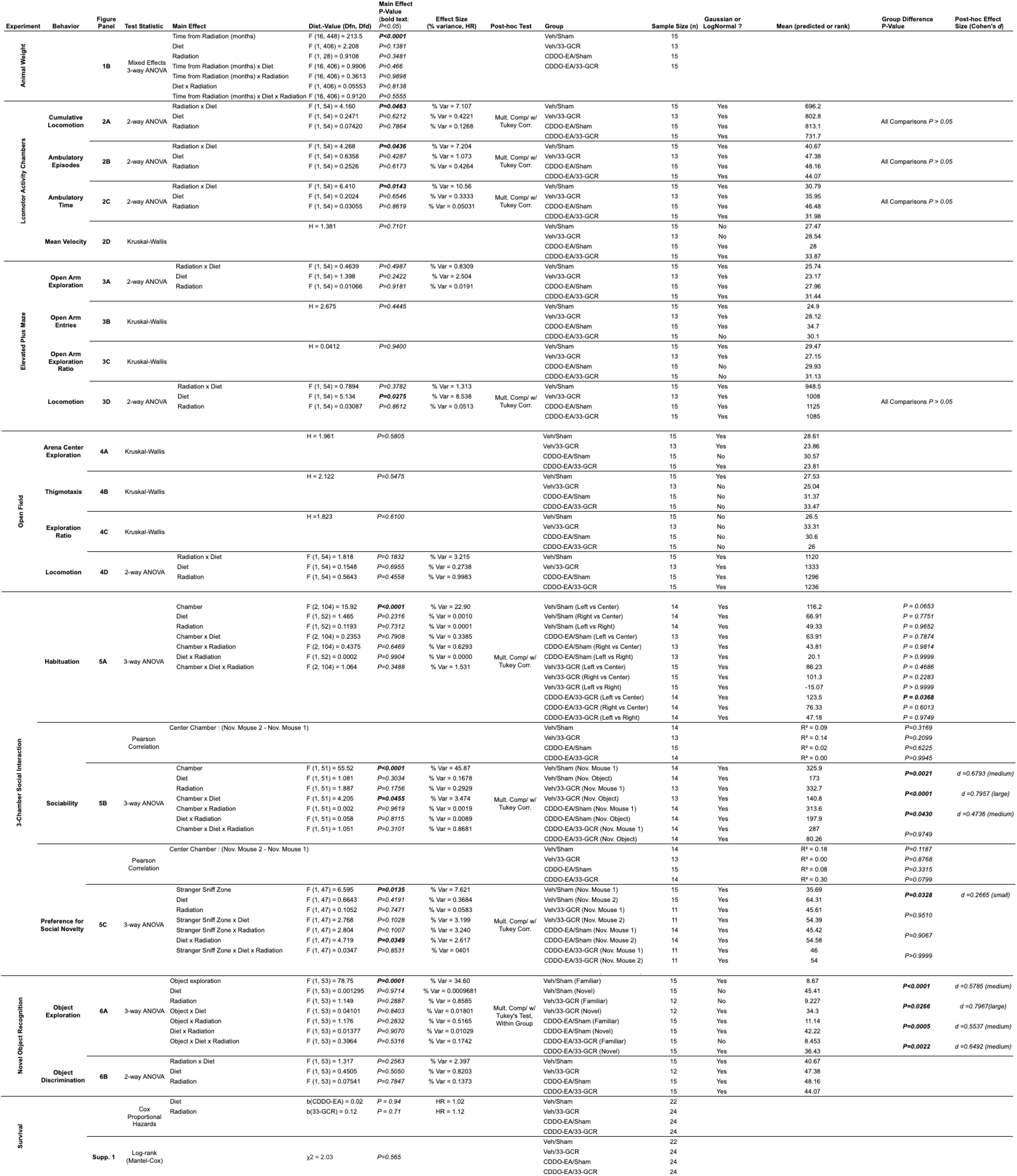
(Statistical Analyses)

**Supplementary Table 2.** Detailed 33-GCR sequential delivery used in this study. Adapted from Simonsen et al. 2020 [12] and adjusted for a total dose of 75cGy.

| Particle | Energy (MeV/n) | Dose Fraction | Dose (cGy) | LET (KeV/um) | Range in H2O (Cm) |
| --- | --- | --- | --- | --- | --- |
| <sup>1</sup>H | 1000 | 24.71% | 18.5325 | 0.22 | 326.6 |
| <sup>4</sup>He | 1000 | 4.98% | 3.735 | 0.88 | 327.8 |
| <sup>28</sup>Si | 600 | 1.62% | 1.215 | 50.2 | 22.73 |
| <sup>1</sup>H | 20 | 6.08% | 4.56 | 2.59 | 0.43 |
| <sup>1</sup>H | 23 | 1.34% | 1.005 | 2.29 | 0.56 |
| <sup>4</sup>He | 20 | 2.20% | 1.65 | 10.34 | 0.43 |
| <sup>4</sup>He | 23 | 0.42% | 0.315 | 9.14 | 0.57 |
| <sup>48</sup>Ti | 1000 | 0.90% | 0.675 | 109.5 | 32.53 |
| <sup>4</sup>He | 27 | 0.44% | 0.33 | 8.06 | 0.75 |
| <sup>4</sup>He | 32 | 0.46% | 0.345 | 7.12 | 0.99 |
| <sup>1</sup>H | 27 | 1.48% | 1.11 | 2.02 | 0.75 |
| <sup>1</sup>H | 32 | 1.60% | 1.2 | 1.79 | 0.98 |
| <sup>1</sup>H | 37 | 1.74% | 1.305 | 1.58 | 1.3 |
| <sup>1</sup>H | 43 | 1.86% | 1.395 | 1.39 | 1.72 |
| <sup>4</sup>He | 37 | 0.50% | 0.375 | 6.29 | 1.31 |
| <sup>4</sup>He | 43 | 0.52% | 0.39 | 5.56 | 1.73 |
| <sup>16</sup>O | 350 | 3.08% | 2.31 | 20.8 | 16.95 |
| <sup>4</sup>He | 50 | 0.54% | 0.405 | 4.92 | 2.28 |
| <sup>4</sup>He | 59 | 0.54% | 0.405 | 4.36 | 3.01 |
| <sup>1</sup>H | 50 | 2.00% | 1.5 | 1.23 | 2.26 |
| <sup>1</sup>H | 59 | 2.12% | 1.59 | 1.09 | 2.99 |
| <sup>1</sup>H | 69 | 2.22% | 1.665 | 0.97 | 3.95 |
| <sup>1</sup>H | 80 | 2.24% | 1.68 | 0.86 | 5.2 |
| <sup>4</sup>He | 69 | 0.54% | 0.405 | 3.86 | 3.97 |
| <sup>4</sup>He | 80 | 0.54% | 0.405 | 3.43 | 5.23 |
| <sup>12</sup>C | 1000 | 2.34% | 1.755 | 7.95 | 110.13 |
| <sup>4</sup>He | 100 | 1.22% | 0.915 | 2.9 | 7.81 |
| <sup>1</sup>H | 100 | 5.44% | 4.08 | 0.73 | 7.76 |
| <sup>1</sup>H | 150 | 7.00% | 5.25 | 0.54 | 15.9 |
| <sup>4</sup>He | 150 | 1.50% | 1.125 | 2.17 | 16 |
| <sup>56</sup>Fe | 600 | 0.82% | 0.615 | 175.1 | 13.09 |
| <sup>4</sup>He | 250 | 3.28% | 2.46 | 1.56 | 38.3 |
| <sup>1</sup>H | 250 | 13.77% | 10.3275 | 0.39 | 38.1 |
| **Total** |  |  | **75** |  |  |

**Supplemental Figure 1.**
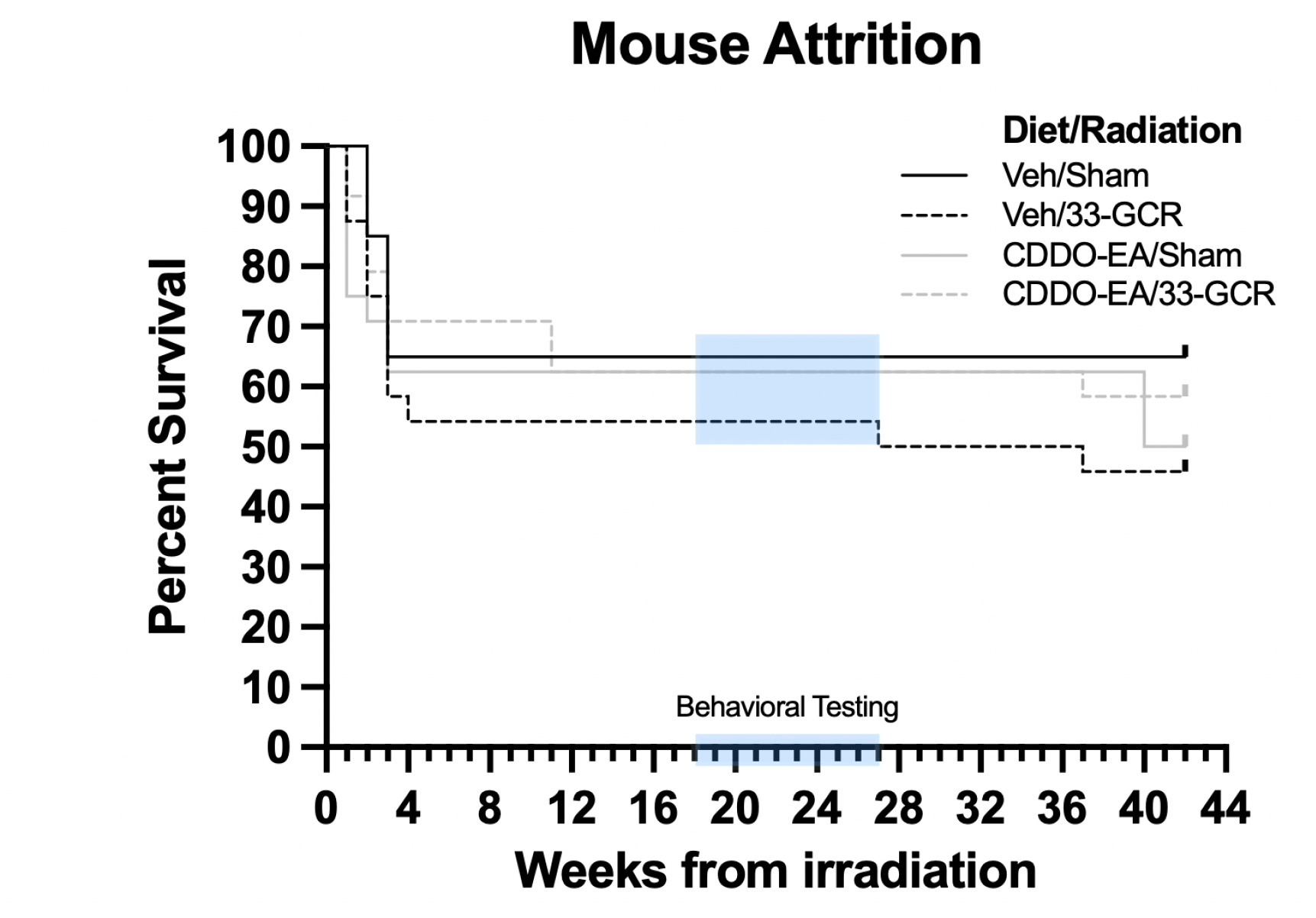
Mouse attrition among groups was not significantly different throughout the study. A Kaplan-Meier survival curve of mice throughout the duration of the study with the behavioral testing period (shaded light blue regions on X axis and on lines). Mice that were singly-housed at the recommendation of veterinarians were counted as losses. Hazard ratio analysis revealed a non-significant non-significant increase in mortality due to CDDO-EA and 33-GCR; therefore, there is negligible hazard due to either Diet or Radiation. Despite a visually lower percent survival in Veh/33-GCR mice vs. the other three groups, Mantel-Cox analysis revealed that survival curves among groups are not significantly different. *n* = 22-24 per group. Details on statistics provided in **Supp. Table 1**.

**Supplemental Figure 2.**
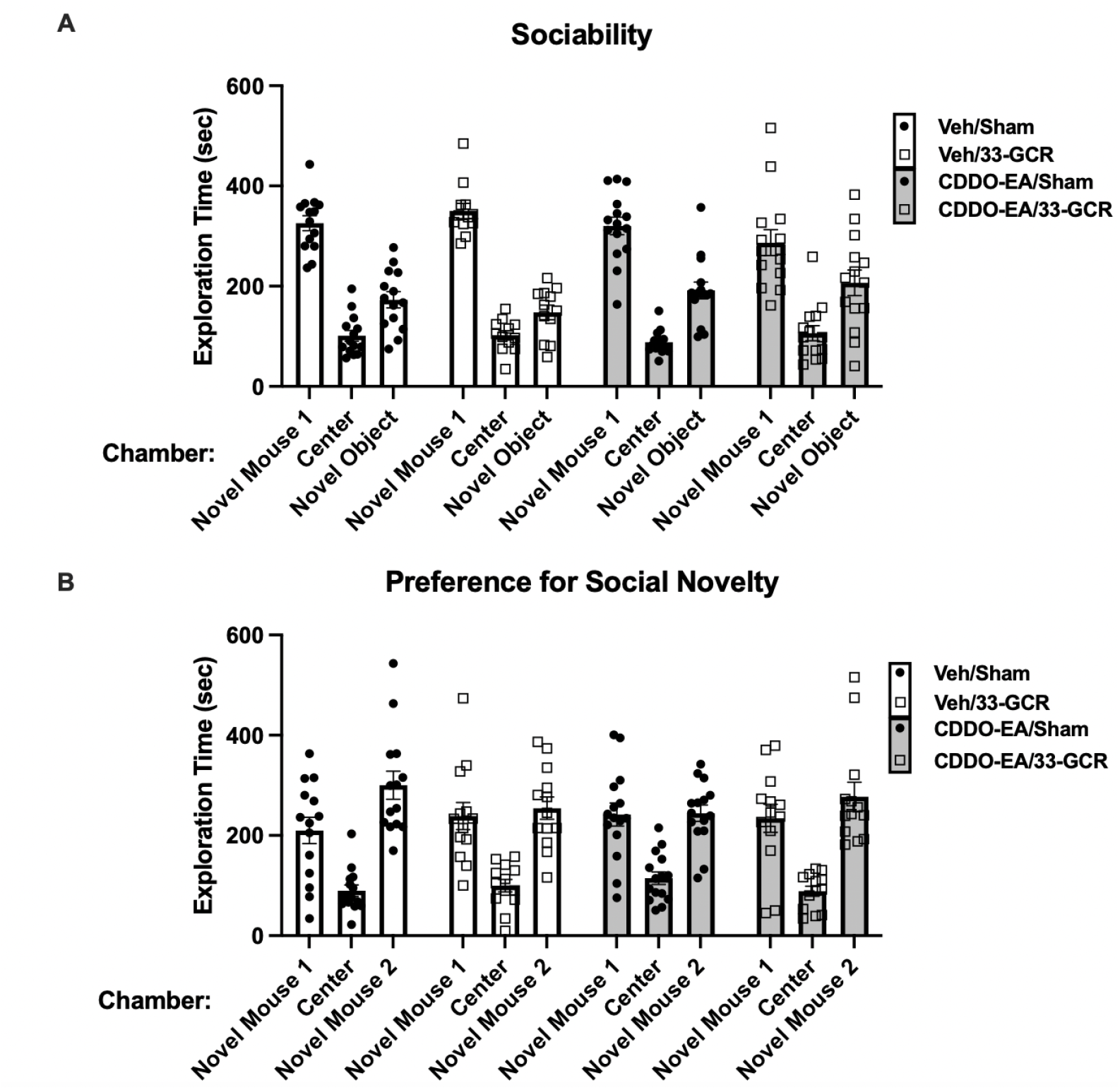
Chamber exploration times for the 3-Chamber Social Interaction Test. Chamber exploration times for the (**A**) Sociability and (**B**) Preference for Social Novelty trials of the 3-CSI test. *n* = 11-15 per group.

**Supplementary Figure 3.**
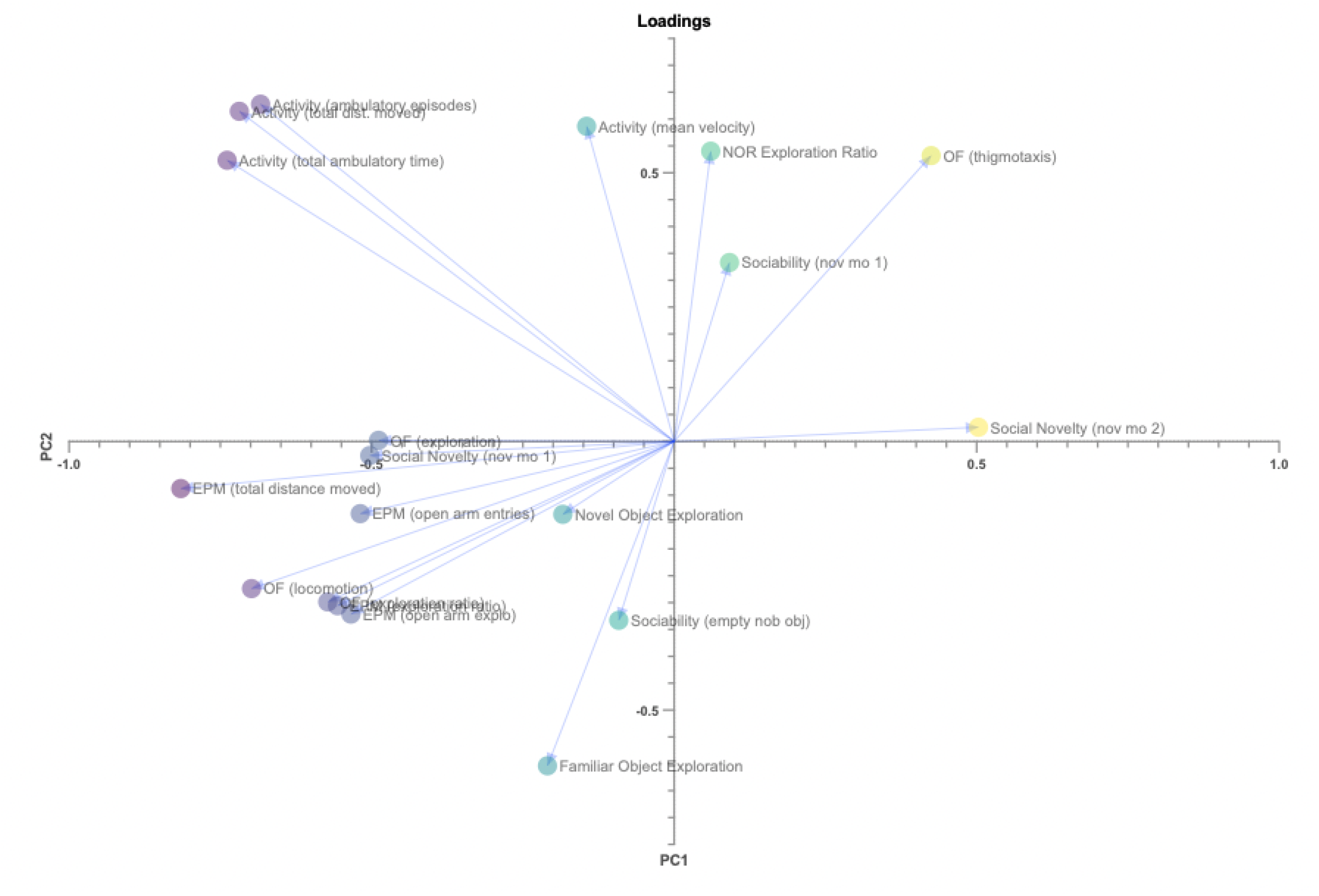
Principal Components Loadings Plot containing each behavioral measure as loadings. Several clusters relating to Anxiety-like, Gross Locomotor Activity, and Exploration behaviors are evident along the PC1 axis, with exploration of novel mouse 2 during the third trial of the 3-chamber social interaction assay as being the behavior with the most isolated variance among all groups in PC1 and PC2.

